# Mapping the unknown: The spatially correlated multi-armed bandit

**DOI:** 10.1101/106286

**Authors:** Charley M. Wu, Eric Schulz, Maarten Speekenbrink, Jonathan D. Nelson, Björn Meder

## Abstract

We introduce the *spatially correlated multi-armed bandit* as a task coupling function learning with the exploration-exploitation trade-off. Participants interacted with bi-variate reward functions on a two-dimensional grid, with the goal of either gaining the largest average score or finding the largest payoff. By providing an opportunity to learn the underlying reward function through spatial correlations, we model to what extent people form beliefs about unexplored payoffs and how that guides search behavior. Participants adapted to assigned payoff conditions, performed better in smooth than in rough environments, and—surprisingly—sometimes performed equally well in short as in long search horizons. Our modeling results indicate a preference for local search options, which when accounted for, still suggests participants were best-described as forming local inferences about unexplored regions, combined with a search strategy that directly traded off between exploiting high expected rewards and exploring to reduce uncertainty about the spatial structure of rewards.

## Introduction

Modern humans descend from capable foragers and hunters, who have migrated and survived in almost every environment on Earth. Our ancestors were able to adaptively learn the distribution of resources in new environments and make good decisions about where to search, balancing the dual goals of *exploring* to acquire new information and *exploiting* existing knowledge for immediate gains. What strategies do humans use to search for resources in unknown environments?

We present a new framework for studying human search behavior using a spatially correlated multi-armed bandit task, where nearby arms (i.e., search options) have correlated rewards. Spatial correlations provide an opportunity to learn about the underlying reward function, extending the traditional reinforcement learning paradigm (Sutton & Barto, 1998) to allow for generalization of learned rewards to unobserved actions using spatial context. We compare search behavior across different payoff conditions, search horizons, and types of environments, finding that participants adapt to their environment, tend to perform very local inferences about unexplored regions and choose arms based on a trade-off between expectations and their attached uncertainties

## Spatially Correlated Multi-Armed Bandits

We adapt the multi-armed bandit (MAB) setting by adding spatial correlation to rewards and placing the arms in a two-dimensional grid (Fig. 1). Each tile represents a playable arm of the bandit, which are initially blank and display the numerical reward value (along with a color aid) after an arm has been chosen. Traditionally, the goal in an MAB task is to maximize cumulative payoffs by sequentially choosing one of the *N*-arms of the bandit that stochastically generate rewards (Steyvers, Lee, & Wagenmakers, 2009), with learning happening independently for each arm (i.e., reinforcement learning). In our case, because proximate arms generate similar rewards, there is the opportunity to form inductive beliefs about unobserved rewards (i.e., function learning). This allows us to study how people generate beliefs about unobserved rewards and how this influences their search behavior.

**Figure 1:**
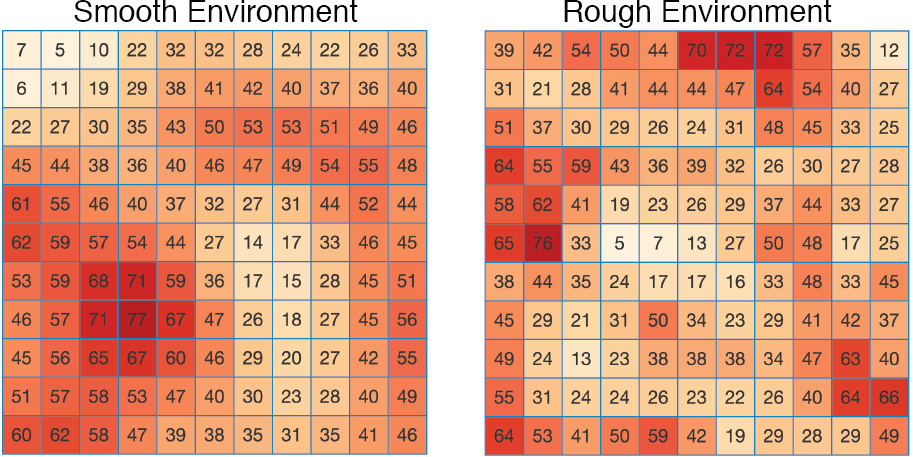
Examples of the underlying reward functions for the two classes of environments.

The spatially correlated MAB is related to the optimal foraging context (Krebs, Kacelnik, & Taylor, 1978), whereby a forager is not only guided by the search for resources, but also by the need to acquire information about the distribution of resources in the environment (Schulz, Huys, Bach, Speeken-brink, & Krause, 2016). This creates a natural trade-off between exploration and exploitation (March, 1991), where an effective search policy needs to adequately balance exploring areas with higher uncertainty, while also exploiting existing information to obtain rewards. One key difference in our task is that the decision-maker must determine *where* to search, and not only whether to stay or to leave a patch.

## Modeling Adaptive Search Behavior

We consider various computational models for describing human behavior, which all make sequential predictions about where people are likely to search. We present both simple strategies without an explicit representation of the environment, along with more complex function generalization models representing the task as a combination of (i) a function learning model and (ii) a decision strategy. We use a form of Gaussian Process regression as a flexible and universal function learning model, which forms inferential beliefs about the underlying reward function, conditioned on previous observations of rewards. Decision strategies are used to transform beliefs into predictions about where to search next. The recovered parameter estimates of our models describe the extent to which people make spatial inferences and how they trade off between exploration and exploitation.

### Simple Strategies

#### Local search

While simple, a tendency to stay local to the previous search decision—regardless of outcome—has been observed in many different contexts, such as semantic foraging (Hills, Jones, & Todd, 2012), causal learning (Bramley, Dayan, Griffiths, & Lagnado, 2017), and eye movements (Hoppe & Rothkopf, 2016). We use inverse Manhattan distance (IMD) to quantify locality:

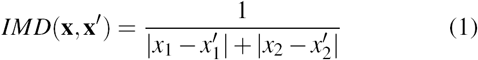

which compares the location of two arms **x** and **x**′, where *x*_1_ and *x*_2_ are the grid coordinates. For the special case where **x** = **x ′**, we set *IMD*(**x**,**x ′)** = 1. At each time *t*, we compute the IMD for each arm based on the choice at **x**_*t-*1_, and then use a softmax function (Eq. 11) to transform locality into choice probabilities, such that arms closer to the previous search decision have a higher probability of being chosen.

#### Win-stay lose-shift

We also consider a form of the winstay lose-shift (WSLS) heuristic (Herbert, 1952), where a *win* is defined as finding a payoff with a higher or equal value than the previous best. When the decision-maker “wins”, we assume that any tile with a Manhattan distance ≤ 1 is chosen (i.e., a repeat or any of the four cardinal neighbors) with equal probability. *Losing* is defined as the failure to improve, and results in choosing any unrevealed tile with equal probability.

### Function Generalization Models

We use a combination of (i) Gaussian Process (*𝒢 𝒫*) regression as a model of how people form beliefs about the underlying reward function conditioned on previous observations (Lucas, Griffiths, Williams, & Kalish, 2015), and (ii) a decision strategy that transforms beliefs into predictions about where a participant will sample next. This approach has recently been applied to human behavior in contextual multi-armed bandits (Schulz, Konstantinidis, & Speekenbrink, 2016) and is the only known computational algorithm to have any guarantees in a bandit setting (i.e., bounded regret; Srinivas, Krause, Kakade, & Seeger, 2010).

#### Gaussian process learning

A *𝒢 𝒫* defines a distribution *P*(*f*) over possible functions *f* (**x**) that map inputs **x** to output *y*, in our case, grid location to reward. A *𝒢 𝒫* is completely defined by a mean *μ*(**x**) and a kernel function, *k*(**x**, **x***′*):

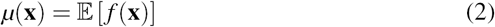

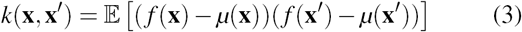

Here, we fix the prior mean to the median value of payoffs, *μ*(**x**) = 50 and use a radial basis function kernel (Eq. 7). Suppose we have collected observations **y**_*T*_ = [*y*_1_*, y*_2_*, …, y*_*T*_]^*T*^ at inputs **X**_*T*_ = *{***x**_1_*, …,* **x**_*T*_}, and assume

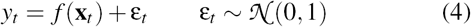

Given a *𝒢 𝒫* prior on functions *f* (**x**) ∼*𝒢 𝒫* (*μ*(**x**)*, k*(**x**, **x** *′*)), the posterior distribution over *f* (**x**_*T*_) given inputs **X**_*T*_ is also a *𝒢 𝒫* with the following mean and covariance:

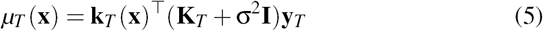

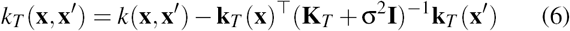

where **k**_*T*_ (**x**) = [*k*(**x**_1_, **x**)*, …, k*(**x**_*T*_, **x**)]^T^ and **K**_*T*_ is the positive definite kernel matrix [*k*(**x**_*i*_, **x** _*j*_)]_*i,* *j*=1*,…,T*_. This posterior distribution is used to derive normally distributed predictions about the rewards for each arm of the bandit (Fig. 2).

**Figure 2:**
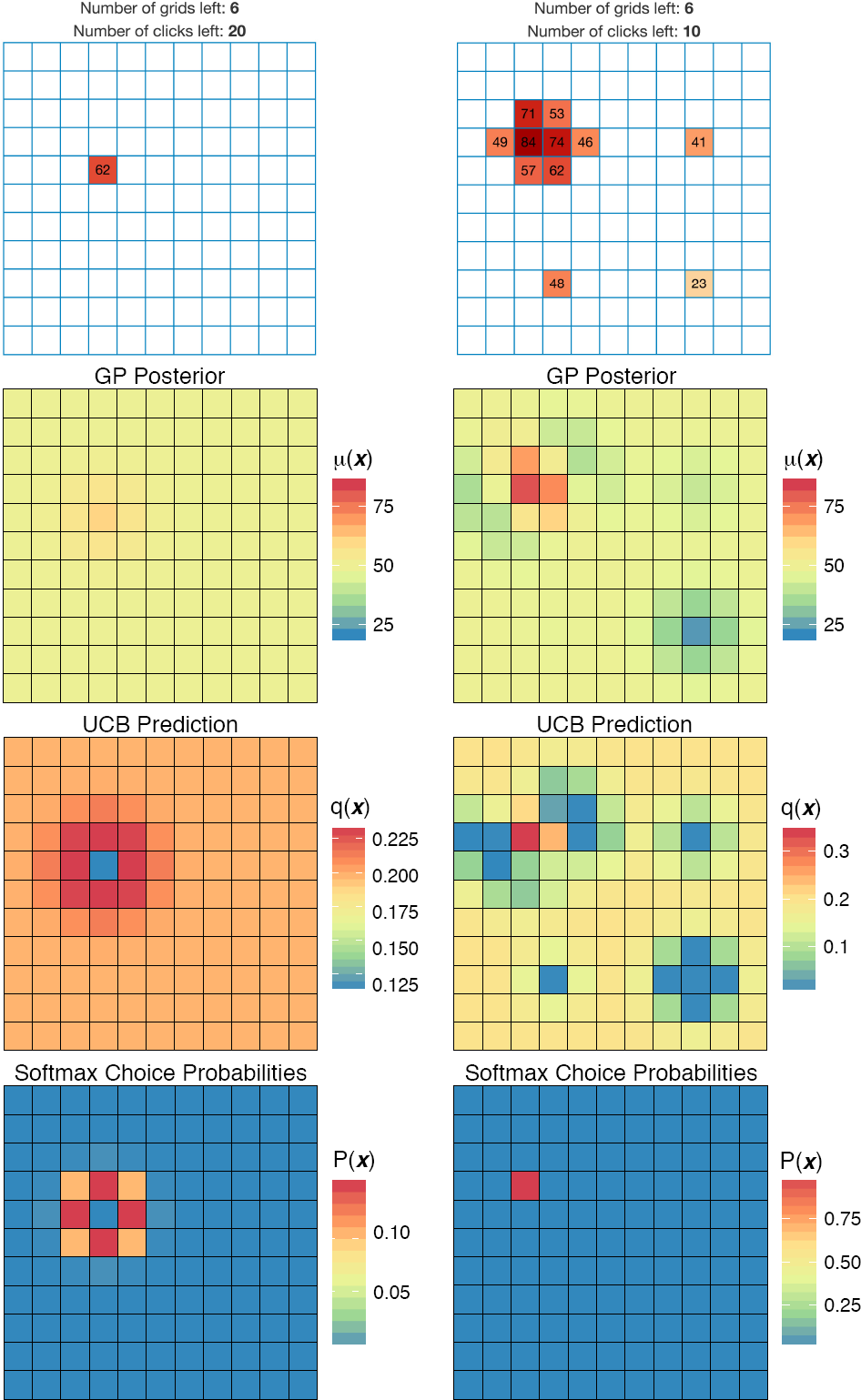
Modeling human performance. Column left represents the initial state of the task and column right is after 10 clicks. Top row: screenshots from the experiment. 2nd row: posterior predictions of expected reward *μ*(**x**), from a *𝒢 𝒫* with an RBF kernel (not shown: the estimated variance). 3rd row: the values of each tile *q*(**x**) using the UCB acquisition function. Bottom row: the softmax prediction surface transforming the UCB values into choice probabilities.

The kernel function *k*(**x**, **x***′*) encodes prior assumptions about the underlying function. We use the *radial basis function* (RBF) kernel

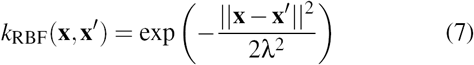

which is a universal function learner and assumes infinitely smooth functions (i.e., correlations between two points **x** and **x** *′* slowly decay as an exponential function of their distance). The RBF kernel uses λ (length-scale) as a free parameter, which determines how far correlations extend: larger values of λ result in longer spatial correlations, whereas λ→ 0^+^ assumes complete independence of spatial information. We use recovered parameter estimates of λ to learn about the extent to which humans make inferences about unobserved rewards.

#### Decision strategies

The *𝒢 𝒫* learning model generates normally distributed predictions about the expectation *μ*(**x**) and the uncertainty σ(**x**) for each arm, which are available to the decision strategies^1^ for evaluating the quality, *q*(**x**), and ultimately making a prediction about where to sample next.

The *Variance Greedy* (VG) strategy values an arm using only the estimated uncertainty

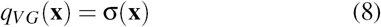

and is an efficient step-wise (greedy) approximation of *information gain* (Srinivas et al., 2010), which seeks to learn the global reward function as rapidly as possible. VG achieves at least a constant fraction of the optimal information gain value (Krause & Guestrin, 2005); however, it fails to adequately trade-off between exploration and exploitation, because effort is wasted exploring the function where *f* (**x**) is small.

The *Mean Greedy* (MG) strategy is also step-wise greedy, valuing arms using only the estimated mean reward

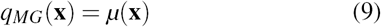

although this strategy carries no known guarantees and is prone to getting stuck in local optima.

*Upper confidence bound sampling* (UCB) combines the VG and MG strategies

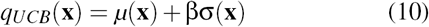

where the exploration factor β determines how the reduction of uncertainty trades off against exploiting high expected rewards. This is sometimes referred to as optimistic “sampling with confidence” as it inflates expectations with respect to the upper confidence bounds (Srinivas et al., 2010), creating a natural balance between exploration and exploitation.

### Choice Probabilities

For all models, we use a softmax function (Fig. 2 bottom row) to convert the value of an option *q*(**x**) into a choice probability

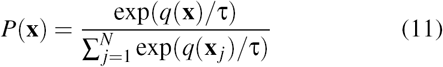

where τ is the temperature parameter. As τ → 0 the highest-value arm is chosen with a probability of 1 (i.e., argmax), and when τ → ∞, all options are equally likely, with predictions converging to random choice. We use τ as a free parameter, where lower estimates can be interpreted as more precise predictions about choice behavior.

## Experiment

We present a bi-variate MAB problem with spatially correlated rewards. The problem space was represented by a two-dimensional grid, measuring 11 × 11, resulting in 121 unique tiles in total. Participants could click to reveal unexplored tiles or re-click previously uncovered tiles to exploit known rewards (see Fig. 2 top row for screenshots).

### Methods

#### Participants

We recruited 80 participants from Amazon Mechanical Turk (25 Female; mean age ± SD 32 ± 9). Each participant was paid a participation fee of $0.50 and a performance contingent bonus up to $1.50. Subjects earned on average $1.64 *±* 0.20 and spent 8 *±* 4 minutes on the task.

#### Design

We used a 2 × 2 between subject design, where participants were randomly assigned to one of two different pay-off structures (*Average Reward* vs. *Maximum Reward*) and one of two different classes of environments (*Smooth* vs. *Rough*). Each grid represented a bi-variate function, with each observation including normally distributed noise, ε∼ *𝒩* (0, 1). The task was presented over 8 blocks on different grid worlds drawn from the same class of environments. In each block, participants had either a *Short* (20 clicks) or *Long* (40 clicks) search horizon to interact with the grid. The search horizon alternated between blocks (within subject), with initial horizon length counterbalanced between subjects. Per block, observations were scaled to a uniformly sampled maximum value in the range of 65 to 85, so that the value of the global optima could not be easily guessed (e.g., a value of 100).

#### Materials and procedure

Before starting, participants were shown four fully revealed grids in order to familiarize themselves with the task. Example environments were drawn from the same class of environments assigned to the participant (Smooth or Rough) and underwent the same random scaling of observations. Additionally, three comprehension questions were used to ensure full understanding of the task.

At the beginning of each of the 8 blocks, one random tile was revealed and participants could use their mouse to click any of the 121 tiles in the grid until the search horizon was exhausted, including re-clicking previously revealed tiles. Clicking an unrevealed tile displayed the numerical value of the reward along with a corresponding color aid, where darker colors indicated higher point values (Fig. 1). Previously revealed tiles could also be re-clicked, although there were variations in the observed value due to noise. For repeat clicks, the most recent observation was displayed numerically, while hovering over the tile would display the entire history of observations. The color of the tile corresponded to the mean of all previous observations.

#### Payoff conditions

We compared performance under two different payoff conditions, requiring either a balance between exploration and exploitation (*Average Reward*) or a pure exploration context (*Maximum Reward*). Previous work has shown that people can adapt (sometimes with great difficulty) to different payoff conditions in information acquisition tasks (Meder & Nelson, 2012).

In each payoff condition, participants received a performance contingent bonus of up to $1.50. *Average Reward* participants were told to “gain as many points as possible across all 8 grids” and were given a bonus based on the average value of all clicks as a fraction of the global optima, 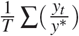 where *y*^*\**^ is the global optimum. *Maximum Reward* participants were told to “learn where the largest reward is” and were giving a bonus using the ratio of the highest observed reward to the global optimum, 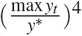, taken to the power of 4 to exaggerate differences in the upper range of performance and for parity in expected earnings across payoff conditions. All 8 blocks were weighted equally, using noisy but unscaled observations to assign a bonus of up to $1.50. Subjects were informed in dollars about the bonus earned at the end of each block.

#### Smoothness of the environment

We used two different classes of environments, corresponding to different levels of smoothness (Fig. 1). All environments were sampled from a *𝒢 𝒫* prior parameterized with a RBF kernel, where the lengthscale parameter (λ) determines the rate at which the correlations of rewards decay over distance. We sampled 20 *Smooth* environments using λ = 2 and 20 *Rough* environments using λ = 1. Subjects performed the task on 8 grids randomly drawn (without replacement) from their assigned class of environments, while the four fully revealed environments used to familiarize subjects with the task were drawn (without replacement) from the remaining 12 environments.

#### Search horizons

The length of the search horizon influences the value of information learned about the environment, with respect to the assigned payoff condition. Longer horizons provide more opportunities for exploiting acquired information, thereby making early exploration more valuable. We chose two horizon lengths (Short= 20 and Long= 40) that were fewer than the total number of tiles on the grid (121), and varied within subject (alternating between blocks).

## Results

Figure 3 shows task performance. In all conditions, performance improved as a function of the trial number (i.e., with each additional click), as measured by both the overall correlation between average reward and trial number (*r* = .32, *p* = .04) and between the maximum observed reward and trial number (*r* = .83, *p* < .001). There were no learning effects across blocks (i.e., over successive grids), indicated by a lack of correlation between average reward and block number (*r* = .19, *p* = .65), or between maximum reward and block number (*r* = -.37, *p* = .36). Performance improved as more information was revealed (i.e., over trials), but not over additional blocks of identically parameterized environments.

**Figure 3:**
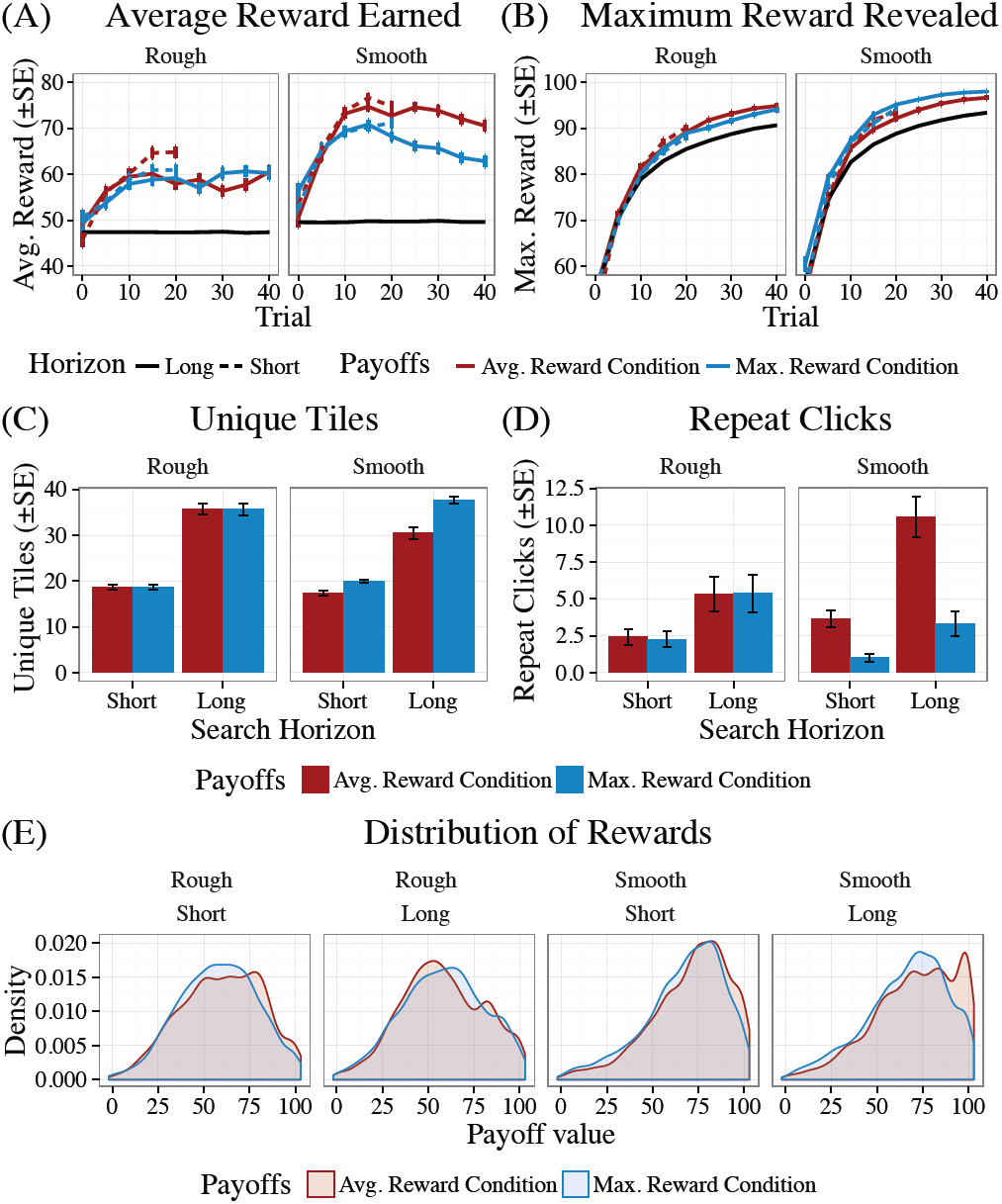
Overview of task performance. (A) Average reward earned and (B) maximum reward revealed, where colors correspond to pay-off condition and line-types to horizon length. Black lines show simulated performance of a random model averaged over 10,000 randomly sampled grids. (C) The average number of unique tiles clicked in each block and (D) the average number of repeat clicks in each block. (E) The distribution of rewards earned during each block, grouped first by environment type and then by horizon length.

### Payoff conditions

Payoff conditions influenced search behavior, with participants in the Maximum Reward condition displaying more variance in the locations sampled (*t*(78) = -2.48, *p* = .02). There were some differences in the number of unique tiles revealed (Fig. 3C) and the number of repeat clicks across the payoff conditions (Fig. 3D), although the effect size is largest for smooth environments given long search horizons. However, these behavioral differences did not manifest in terms of performance, with no systematic differences across payoff conditions in terms of the average reward obtained *t*(78) = 1.32, *p* = .2) or in the maximum revealed reward (*t*(78) = .001, *p* = .99).

### Environment and horizon

Independent of the payoff condition, participants assigned to Smooth environments achieved higher average rewards (*t*(78) = 6.55, *p <* .001) and higher maximum rewards (*t*(78) = 5.45, *p < .*001), than those assigned to the Rough environments (Fig. 3E), suggesting that stronger correlations of payoffs make the task easier. Interestingly, longer horizons did not lead to better overall performance in the Average Reward condition (*t*(80) = .34, *p* = .73), although participants given longer horizons found larger maximum rewards for all payoffs and environment conditions (*t*(158) = 7.62, *p < .*001). There may be a less-is-more-effect, with longer horizons leading to over-exploration, given the goal of maximizing average rewards.

### Model Comparison

We describe each model’s ability to predict participant behavior using leave-one-block-out cross validation. For each participant, we analyzed the four short and the four long horizon blocks separately. Cross-validation was performed by holding out a single block as a test set, and fitting the model parameters using a maximum likelihood estimate (MLE) on the remaining three blocks. Iterating through each of the four hold-out blocks, for both short and long horizons, we calculated a model’s out-of-sample log loss (i.e., test set prediction accuracy) and then summed up the results over all blocks. We use McFadden’s *R*^2^ values (McFadden, 1974) to compare the out-of-sample log loss for each model to that of a random model (Fig. 4), where *R*^2^ = 0 indicates chance performance and *R*^2^ = 1 is a perfect model.

**Figure 4:**
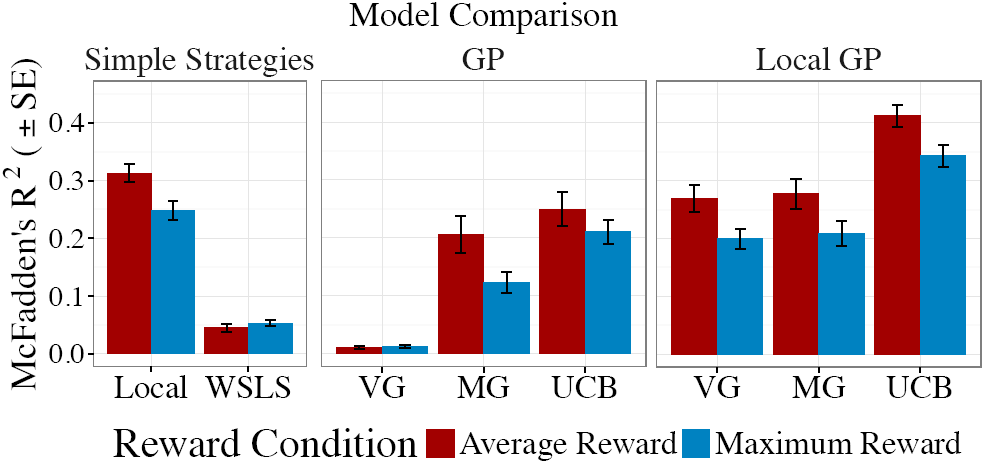
Model Comparison. The height of the bars show the group mean and error bars indicate standard error. McFadden’s *R*^2^ is a goodness of fit measure comparing each model *ℳ*_*k*_ to a random model *ℳ*_rand_. Using the out-of-sample log loss for each model, 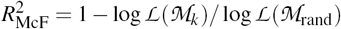

A large amount of the variance in participant behavior is explained by local search (*R*^2^ = .28; all conditions); however, locality alone fails to achieve similar task performance as humans, with performance almost identical to random in terms of average reward and worse than random in maximum reward (Fig. 5). WSLS by comparison, was a poor approximation of search behavior (*R*^2^ = .05), and was excluded from the model performance comparison.

**Figure 5:**
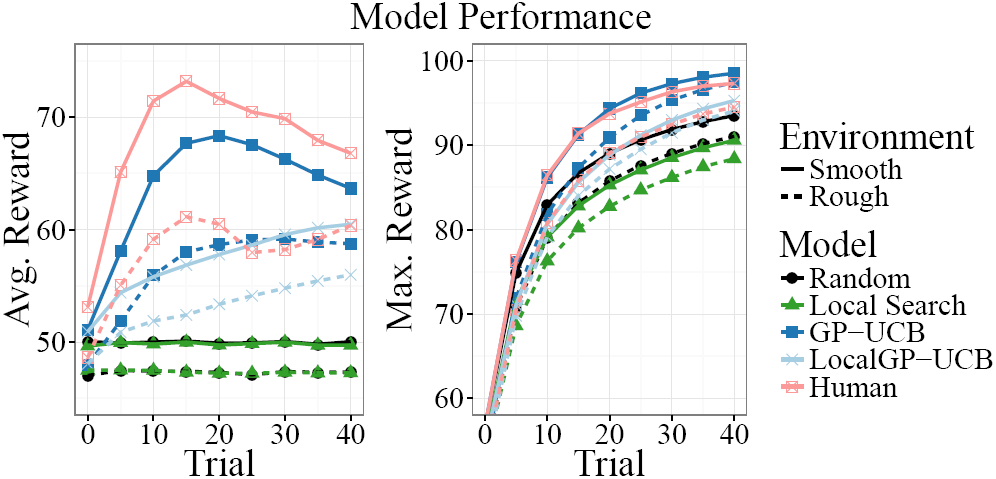
Comparison of simulated model performance over 10,000 replications, where parameters were sampled from the crossvalidated MLEs of the subject population. Human results are averaged across payoff conditions and horizon length.

Among the GP models, UCB performed best (*R*^2^ = .23), with MG showing comparable results (*R*^2^ = .17) and VG performing poorly (*R*^2^ = .01). Interestingly, the performance of the GP-UCB model was remarkably similar to human subjects in terms of both average and maximum reward (Fig. 5). Both humans and the GP-UCB model explore beyond what is adaptive in the average reward context as evidenced by the peak around *t* = 15, continuing to explore after most high-value rewards have been revealed and thus failing to consistently improve average rewards^2^.

To harmonize the different aspects of human behavior captured by local search and by the GP-UCB model, we added a local variant of each GP model (*Local GP*), which weighs the *q*(**x**) for each arm by the inverse Manhattan distance to the previous choice, *q*_*Local*_ (**x**_*t*_) = *q*(**x**_*t*_) *IMD*(**x**_*t*_, **x**_*t-*1_). Adding locality to the GP models only improved prediction accuracy (Fig. 4 right), with the Local GP-UCB model having the highest overall out-of-sample prediction accuracy (*R*^2^ = .38).

Overall, the modeling results show that humans display a preference for local search, but that locality alone fails to achieve comparable performance levels. The best model (Local GP-UCB) incorporated this tendency for local search into a computational model that combines function learning with a decision strategy explicitly trading off between both high expected rewards and high uncertainty.

### Parameter Estimation

Figure 6 shows the cross-validated parameter estimates of the best predicting Local GP-UCB model. The estimates indicate subjects systematically under-estimated the smoothness of the underlying environments, with λ values lower than the true underlying function (λ_*Smooth*_ = 2, λ_*Rough*_ = 1), for both Rough environments (*t*(36) = -4.80, *p < .*001) and Smooth environments (*t*(42) = -18.33, *p < .*001), using the median parameter estimate for each subject. Participants not only had a tendency towards selecting local search options, but also made local inferences about the correlation of rewards.

**Figure 6:**
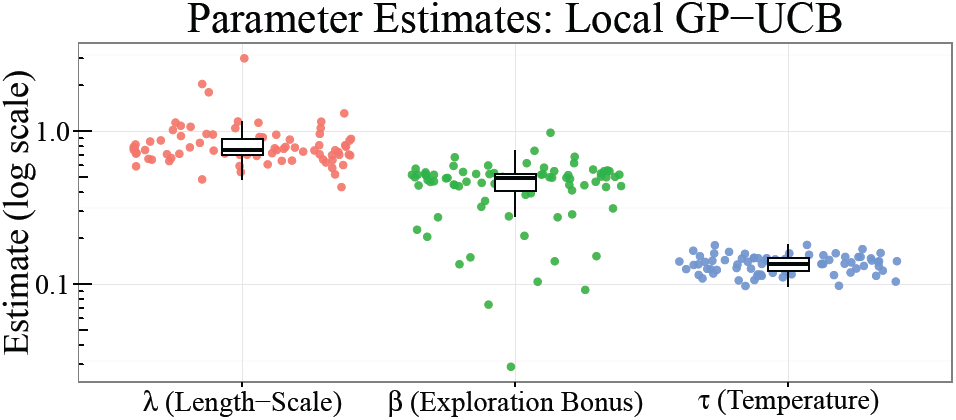
Cross-validated parameter estimates for the Local GP-UCB model, showing the median estimate for each participant.

All participants valued the reduction of uncertainty (β *>* 0), with long horizons often yielding larger β estimates than short horizons (51 out of 80 subjects; *t*(79) = -2.02, *p* =.047)^3^. There were no differences between payoff conditions (*t*(78) = -1.65, *p* = .1) or environments (*t*(78) = .5, *p > .*1).

Subjects in the average reward condition yielded smaller estimates of the softmax temperature parameter (τ) than those in the maximum reward condition (*t*(78) = -2.66, *p* = .009), This is consistent with almost all models making better predictions for average reward than for maximum reward subjects (Fig. 4), since smaller values of τ indicate more precise predictions. The larger number of unique tiles searched in the maximum reward condition (Fig. 3C) may indicate a more difficult prediction problem.

## General Discussion

The results presented here can be seen as a first step towards uncovering how people search to acquire rewards in the presence of spatial correlations. We have re-cast the multi-armed bandit problem as a framework for studying both function-learning and the exploration-exploitation trade-off by adding spatial correlations to rewards. Within a simple experiment about searching for rewards on a two-dimensional grid, we found that participants adapt to the underlying payoff condition, perform better in smooth than in rough environments, and—surprisingly—sometimes seem to perform as well in short as in long horizon settings.

Our modeling results show a tendency to prioritize local search options, which may indicate the presence of innate search costs (e.g., mouse movements or some additional cognitive processing). Even accounting for this local search behavior, our best predicting model (Local GP-UCB) indicates that people still systematically underestimate the extent of spatial correlation of rewards, preferring instead to make very local inferences about unexplored rewards. Additionally, we also found that search behavior was best predicted by a combination of both high expected reward and high uncertainty, embodied in the UCB decision strategy, which implicitly negotiates the exploration-exploitation trade-off.

Future studies could expand on this work by assessing a more diverse and perhaps combinatorial set of kernel functions (Schulz, Tenenbaum, Duvenaud, Speekenbrink, & Gershman, 2016) or by speeding up GP-inference using approximation methods such as sparse inference (Lawrence, Seeger, & Herbrich, 2003) or more parsimonious neural network representations (Neal, 2012). Indeed, the result that participants formed only very local beliefs about spatial correlations could be used to find heuristic approximations to GP models in the future, which could effectively trade-off a small loss in accuracy for reduced computational complexity.

## Conclusion

We compared both simple strategies and more complex function generalization models in their ability to make out-of-sample predictions about participant sampling behavior. Our modeling results indicate that there may be innate search costs, creating a tendency to prioritize local search options. Furthermore, even accounting for this local search behavior, our best performing model (Local GP-UCB) indicates that people also have a systematic tendency to underestimate the extent of spatial correlation of rewards, preferring instead to make very local inferences about unexplored rewards.

## Acknowledgments

CMW is supported by the International Max Planck Research School on Adapting Behavior in a Fundamentally Uncertain World. ES is supported by the UK Centre for Financial Computing and Analytics. BM was supported by Grant ME 3717/2-2, JDN was supported by Grant NE 1713/1-2 from the Deutsche Forschungsgemeinschaft (DFG; SPP 1516). Code is available at https://github.com/charleywu/gridsearch

## Footnotes

1 We also considered Probability of Improvement and Probability of Maximum Utility (Speekenbrink & Konstantinidis, 2015) as alternate decision strategies, but have omitted them because they failed to reach performance comparable to UCB.

2 Note that the peak in average reward for the GP-UCB is due to the use of human parameter estimates, whereas a GP-UCB model with optimized hyper-parameters and a dynamic β is known to achieve sublinear regret bounds (i.e., monotonically increasing average reward; Srinivas et al., 2010)

3 Because horizon length varied within subjects, we compare the aggregate mean of the cross-validated parameter estimates for β.

